# *Opa1* overexpression protects from early onset *Mpv17*^*-/-*^-related mouse kidney disease

**DOI:** 10.1101/2020.03.18.996561

**Authors:** Marta Luna-Sanchez, Cristiane Benincà, Raffaele Cerutti, Gloria Brea-Calvo, Anna Yeates, Luca Scorrano, Massimo Zeviani, Carlo Viscomi

## Abstract

Moderate overexpression of *Opa1*, encoding a master regulator of mitochondrial cristae morphology, has been shown to improve significantly mitochondrial damage induced by drugs, surgical denervation, or genetically determined OXPHOS defects. However, this approach has been so far demonstrated in a limited number of genetically defective OXPHOS models characterized by specific impairment of a single mitochondrial respiratory chain complex. Here, we investigated the effectiveness of moderate *Opa1* overexpression in the *Mpv17*^*-/-*^ mouse, characterized by profound, multisystem mtDNA depletion. In naïve *Mpv17*^*-/-*^ individuals, whose genetic background was crossed with individuals belonging to the *Opa1^*tg*^* strain, we found a surprising anticipation of severe, progressive focal segmental glomerulosclerosis, previously described in *Mpv17*^*-/-*^ animals as a late-onset clinical feature (after 12-18 months of life). In contrast, kidney failure led *Mpv17*^*-/-*^ individuals from this new “mixed” strain leading to death 8-9 weeks after birth. However, *Mpv17*^*-/-*^::*Opa1*^*tg*^ mice lived much longer than *Mpv17*^*-/-*^ littermates, and developed much later severe proteinuria associated with focal segmental glomerulosclerosis. MtDNA content and OXPHOS activities were significantly higher in *Mpv17*^*-/-*^::*Opa1*^*tg*^ than in *Mpv17*^*-/-*^ kidneys, and similar to WT littermates. Mitochondrial network and cristae ultrastructure were largely preserved in *Mpv17*^*-/-*^::*Opa1*^*tg*^ vs. *Mpv17*^*-/-*^ kidney and isolated podocytes. Mechanistically, the protective effect of *Opa1* overexpression in this model was mediated by a block in apoptosis due to the stabilization of the mitochondrial cristae, consequently increasing the levels of mitochondrial morphology proteins like MFN2 and MIC19 as well as stabilizing ATP synthase oligomers. These results demonstrate that strategies aiming at increasing *Opa1* expression or activity can be an effective aid against mtDNA depletion syndromes.

## Introduction

There is an unmet need for new therapies for mitochondrial diseases. Given the extreme genetic, biochemical and clinical heterogeneity of mitochondrial disorders, an ideal therapeutic approach should be applicable to more than one single clinical entity. Strategies with potential general applicability to several, if not all, mitochondrial diseases have been proposed over the last 10 years by us and others (Viscomi & Zeviani, 2020). These include approaches aiming at (i) increasing mitochondrial biogenesis, (ii) clearing dysfunctional organelles by stimulating mitophagy, and, although more controversially, (iii) through the by-pass of mitochondrial respiratory chain (MRC) complexes by using alternative oxidases, or (iv) by reversing the mitochondrial unfolded stress response (Viscomi & Zeviani, 2020). An additional, promising approach is based on the moderate overexpression *Opa1*, encoding a master regulator of shape and function of mitochondrial cristae. We previously showed that this approach improved the clinical phenotype of the *Ndufs4*^*−/−*^ mouse, which lacks a small subunit of the P-module of the MRC complex I (cI), as well as of the muscle-restricted *Cox15*^*sm/sm*^, which lacks an enzyme essential for the biosynthesis of COX-specific hemeA (Civiletto *et al*, 2015). We proposed that the stabilization of the defective MRC and supercomplexes, due tighter and tidier cristae morphology, mediated the therapeutic effect. In addition, the same approach protected mice from a number of insults leading to altered cristae morphology and decreased OXPHOS proficiency, including denervation, LPS-induced liver damage, and ischemia-reperfusion in heart and brain (Varanita *et al*, 2015). However, the applicability of the *Opa1* overexpression therapy to other genetically determined mitochondrial diseases, such as those due to instability of mtDNA, has not been investigated. Here, we sought to determine if *Opa1* overexpression could improve the molecular phenotype of the *Mpv17* knockout mouse associated with mtDNA depletion, which mimics the molecular features of a severe mitochondrial syndrome in humans due to *Mpv17* mutations. *Mpv17* encodes a small and highly hydrophobic protein embedded in the inner mitochondrial membrane (IMM), whose function is still unknown. In humans, mutations in *MPV17* cause hepatocerebral mtDNA depletion syndrome (OMIM #266810) (Spinazzola *et al*, 2006), including the Navajo neurohepatopathy (Karadimas *et al*, 2006). This syndrome is characterized by early-onset and profound mtDNA depletion in liver, later complicated by neurological failure and, in some cases, peripheral analgesia with corneal scarring. Milder reduction of mtDNA content in skeletal muscle and, occasionally, multiple mtDNA deletions, in addition to mtDNA depletion in liver, have also been reported (Piekutowska-Abramczuk *et al*, 2014; Uusimaa *et al*, 2014). The patients surviving the acute hepatic failure, dominated by hypoglycaemic crises and cirrhotic evolution of the liver parenchyma, develop a progressive ataxia and neurological impairment. The spectrum of disorders due to mutations in *MPV17* has progressively expanded to include (i) juvenile and adult-onset axonal sensorimotor polyneuropathy without hepatocerebral involvement (Baumann *et al*, 2019; Choi *et al*, 2015); (ii) neuropathy and leukoencephalopathy with multiple mtDNA deletions in skeletal muscle (Blakely *et al*, 2012); (iii) adult-onset neurohepatopathy-plus syndrome with multiple deletions of mtDNA in muscle (Garone *et al*, 2012). The common feature of all these clinical syndromes is mtDNA instability, pointing to a role of MPV17 in mtDNA maintenance. However, the function of the *Mpv17* gene product remains unclear, although several hypotheses have been proposed. For instance, MPV17 has been shown to form a non-selective cation channel, affecting mitochondrial membrane potential and ROS production (Antonenkov *et al*, 2015; Reinhold *et al*, 2012). Other data suggested that the lack of MPV17 leads to a substantial decrease in dGTP and dTTP in mouse liver mitochondria, thus facilitating the incorporation of riboGTP, which, in turn, distorts mtDNA and may lead to block of its replication (Dalla Rosa *et al*, 2016). Accordingly, high levels of rGTP were detected in mtDNA from *Mpv17*^*-/-*^ livers (Moss *et al*, 2017). Other data showed that reduced *MPV17* expression was associated with impaired dTMP synthesis without affecting de novo or salvage synthesis of dTMP, suggesting that MPV17 may be involved in maintaining dTMP levels in mitochondria through a still uncharacterized pathway, which may be involved in transporting dTMP or one of its precursors from the cytosol to mitochondria (Alonzo *et al*, 2018). In zebrafish, *Mpv17* has been related to pyrimidine nucleotide metabolism via impairment of dihydroorotate dehydrogenase (Martorano *et al*, 2019). All these observations have been reported in single publications and do need further confirmation. Finally, data in *Saccharomyces cerevisiae* support a role for *Sym1*, the yeast orthologue of *Mpv17*, in the homeostatic control of TCA cycle intermediates (Dallabona *et al*, 2010).

Similar to mutant *MPV17* patients, *Mpv17*^*-/-*^ mice are characterized by profound depletion of mtDNA in liver, and moderate to mild depletion in other organs, such as skeletal muscle, brain and kidney. However, despite the profound liver mtDNA depletion and contrary to the severe liver failure that characterizes the patients, *Mpv17*^*-/-*^ mice survive normally up to one year of age, except an invariant greying of their fur occurring around the sixth month of life. However, after the first year of life, they develop a focal segmental glomerulosclerosis, leading to death by 1.5-2.0 years of age (Viscomi *et al*, 2009). A yet unexplained finding concerning *Mpv17*^*-/-*^ mouse model is the shift of the kidney failure, which led to early death a few weeks after birth in the original strain (Weiher *et al*, 1990; O’Bryan *et al*, 2000), to a much later and slowly progressive condition that was observed and documented in mouse *Mpv17*^*-/-*^ individuals after several generations (Viscomi *et al*, 2009).

## Results

### Early onset kidney disease in *Mpv17*^*-/-*^ mice

In order to generate *Mpv17*^*-/-*^::*Opa1*^*tg*^ double recombinant mice, we mated *Mpv17*^*+/-*^ and *Opa1*^*tg*^ mouse strains. The latter express *Opa1* to slightly higher levels than controls (CTRs) by targeted insertion of an extra copy of *Opa1* into the *Hprt* locus on the X chromosome (Cogliati *et al*, 2013; Civiletto *et al*, 2015; Varanita *et al*, 2015). Surprisingly, we noticed that the first *Mpv17*^*-/-*^ animals (without *Opa1* extra copies) started to die around 8-9 weeks of age (Figure 1A). This was reminiscent of the original phenotype described for this strain, i.e. an early death due to focal segmental glomerulosclerosis. We termed this strain as “new” *Mpv17*^*-/-*^ strain, compared to our later-onset slowly progressive “old” *Mpv17*^*-/-*^ strain. To search for possible modifier genes responsible of the phenotype anticipation, we first analysed two genes previously proposed to modify the severity of the clinical phenotype in some mouse models of mitochondrial dysfunction, i.e. *Prkdc*, encoding for the catalytic subunit of the DNA-dependent protein kinase (DNA-PK) (Papeta et al, JCI 2010), and *Nnt*, encoding the nicotinamide nucleotide transhydrogenase (McManus *et al*, 2019). However, the *Prkdc* mutation described in Balb/c and other inbred strains was not present in our C57Bl/6 colonies, and the *Nnt* mutation, present in the of pure C57Bl/6J strain, did not segregate with the disease (not shown). In order to identify potential genetic modifiers, we also carried out an RNA sequencing (RNAseq) experiment, by analysing the total RNA extracted from the kidneys of four *Mpv17*^*-/-*^ mice obtained after the crossing with the *Opa1*^*tg*^ mice, and showing early-onset kidney failure, vs. four individuals belonging to our original, long surviving, *Mpv17*^*-/-*^ strain, sacrificed at one month of age, i.e. in a pre-symptomatic stage in both strains. The results are summarized in the Volcano plot in Supplementary Figure S1A. We found that 23 genes were significantly varied in the new vs. original strain. Among these, the only mitochondrial ones were the *Slc25a48*, encoding an uncharacterized carrier, which was upregulated in the new strain, and *Upp2*, encoding the uridine phosphorylase 2, which was downregulated in the new strain. Other gene products differentially expressed in the two strains did not localize in mitochondria, and the most differently expressed target was a non-coding RNA, termed AC158975.2, which was downregulated in the new KO strain. These results are also reported in detail in Supplementary Figure S1B.

**Figure 1.**
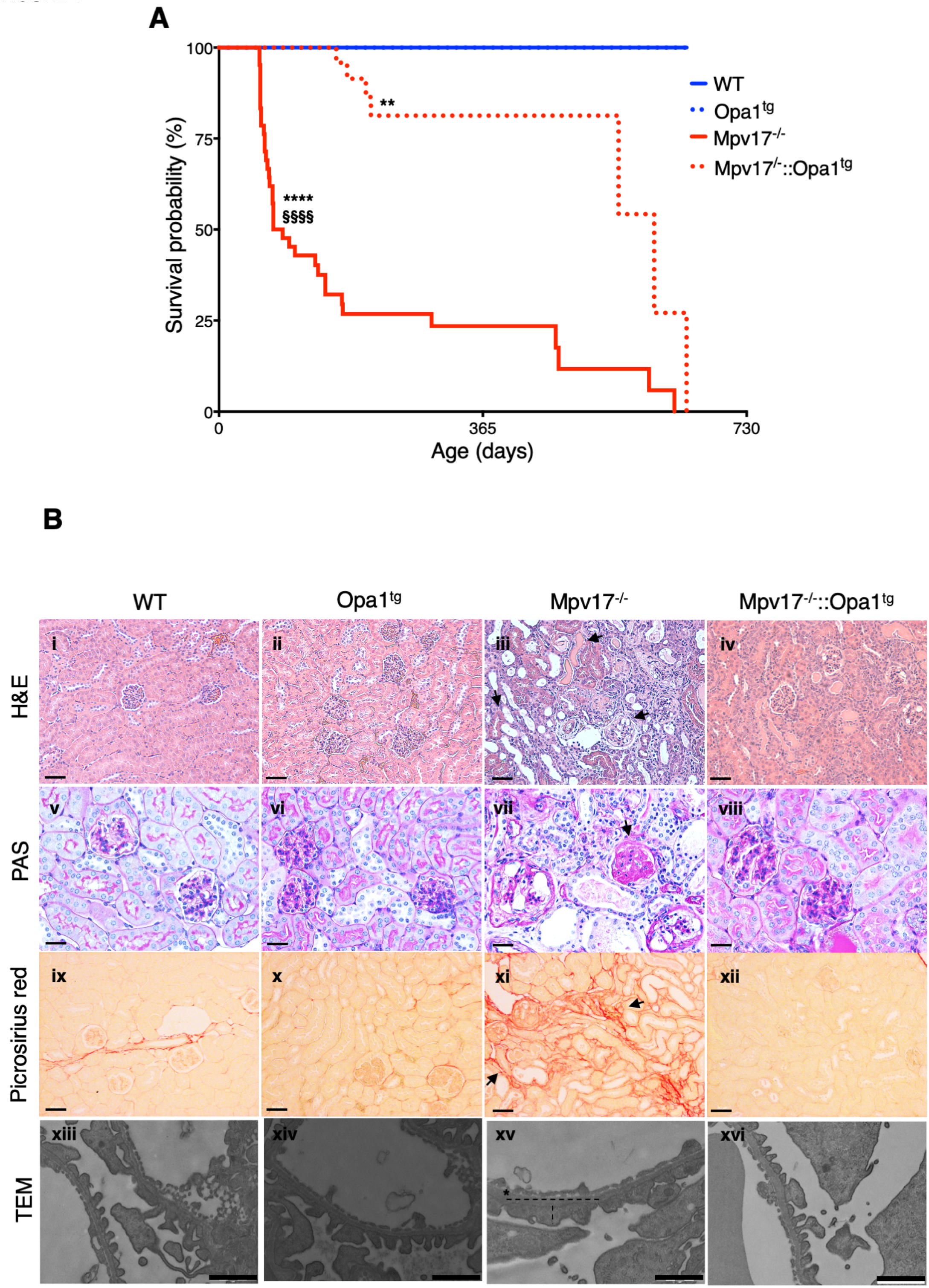
Effects of Opa1 overexpression on survival probability and kidney disease of Mpv17^−/−^ mice. A) Kaplan-Meier analysis of WT, *Opa*^*tg*^, *Mpv17*^*-/-*^ and *Mpv17*^*-/-*^::*Opa1*^*tg*^ mice (n = 30). Kaplan-Meier survival probability. Significance was assessed by the log rank test. Symbols * and **§** represent the significance levels vs. WT and *Mpv17*^*-/-*^, respectively: **** p<0.0001 (*Mpv17*^*-/-*^ vs. WT), ** p=0.0067 (*Mpv17*^*-/-*^::*Opa1*^*tg*^ vs. WT), **§§§§** p<0.0001 (*Mpv17*^*-/-*^::*Opa1*^*tg*^ *vs. Mpv17*^*−/−*^). B) Histological characterization of WT, *Opa1*^*tg*^, *Mpv17*^*-/-*^ and *Mpv17*^*-/-*^::*Opa1*^*tg*^ kidneys. Scale bar: 50μm for all the light microscopy stainings except for PAS for which the bar size is 20μm. TEM shows foot process in kidney of the indicated genotypes. Dashed line indicates height and width of foot process, the asterisk indicates the glomerular basement membrane (GBM). Scale bar: 1 μm.

### *Opa1* moderate overexpression prolongs the lifespan of *Mpv17*^*-/-*^ mice by preventing early onset glomerulosclerosis

To investigate the presence of kidney failure in our strains, we measured proteinuria by using semi-quantitative strips. All tested “new” *Mpv17*^*-/-*^ animals displayed high proteinuria compared to the CTR littermates (n=8/group) (Supplementary Figure S2A). Necroscopic analysis of 8-week old mice revealed that the kidneys were smaller in *Mpv17*^*-/-*^ *vs.* CTR littermates, which included both *Mpv17*^*+/+*^ and *Mpv17*^*+/−*^ individuals (Supplementary Figure 2B). Hematoxylin and eosin (H&E) staining showed the presence of many degenerating glomeruli and some protein casts in the tubuli of *Mpv17*^*-/-*^ (Figure 1B) but neither in the tubuli of CTR nor of *Opa1*^*tg*^ mice (Figure 1B). PAS staining revealed the presence of collagen-related glycoproteins in the glomeruli, with partial obliteration of the capillary loops (Figure 1B). Picrosirius-red staining confirmed the massive presence of fibrotic tissue (Figure 1B). Finally, Transmission Electron Microscopy (TEM) showed profoundly altered podocytes in the glomeruli, with loss of the foot processes (Figure 1B and Supplementary Figure 2C). These data confirmed the presence of an early-onset and severe focal segmental glomerulosclerosis in the “new” *Mpv17*^*-/-*^ mouse strain. However, *Opa1* overexpression had a striking protective effect, so that *Mpv17*^*-/-*^:: *Opa1*^*tg*^ mice lived significantly longer than naïve *Mpv17*^*-/-*^ littermates (median lifespan: 553 vs. 75 days) (Figure 1A), and showed only minor histological alterations (Figure 1B).

### *Opa1* overexpression prevents OXPHOS defects by increasing mtDNA content in kidney

Profound mtDNA depletion in liver and more moderate in kidney is a hallmark of *Mpv17*^*-/-*^ mice (Viscomi *et al*, 2009). We thus investigated whether *Opa1* overexpression was able to increase mtDNA content in these tissues. We found that liver mtDNA content was doubled in *Mpv17*^*-/-*^::*Opa1*^*tg*^ mice compared to “new” *Mpv17*^*-/-*^ animals (10.92±2.33 vs. 4.31±2.28, p<0.01, n=6/genotype), but it was still markedly lower than in WT or *Opa1*^*tg*^ littermates (100±33 and 106±20, respectively p<0.0001, n=6) (Supplementary Figure S3A). Accordingly, the spectrophotometric activities of MRC complexes cI, cIII_2_, and cIV in liver homogenates were no different between *Mpv17*^*-/-*^::*Opa*^*tg*^ and *Mpv17*^*-/-*^ and both were significantly lower than in CTR and *Opa1*^*tg*^ groups (Supplementary Figure S3B), despite *Opa1* overexpression was confirmed in the liver of *Mpv17*^*-/-*^:: *Opa1*^*tg*^ *vs. Mpv17*^*−/−*^ animals (Supplementary Figure S3C).

We then analysed the kidneys, the most affected organ in this “new” *Mpv17*^*-/-*^ model characterized by early-onset disease. First, we confirmed that *Opa1* was overexpressed in *Opa1*^*tg*^ and *Mpv17*^*-/-*^::*Opa1*^*tg*^ compared to WT and *Mpv17*^*-/-*^ by real time PCR (Supplementary Figure S4A, B) and Western-blot immunovisualization (Supplementary Figure S4B). Second, we measured mtDNA content. Similarly to liver, mtDNA content was doubled in *Mpv17*^*-/-*^:: *Opa1*^*tg*^ vs. *Mpv17*^*-/-*^ kidneys (69±13.5 Vs. 33±21.1). The mtDNA amount of double recombinant *Mpv17*^*-/-*^:: *Opa1*^*tg*^ individuals was indeed not significantly different from that of WT and *Opa1*^*tg*^ mice (100.0±13.0 and 90.0±15.3, respectively) (Figure 2A).

**Figure 2.**
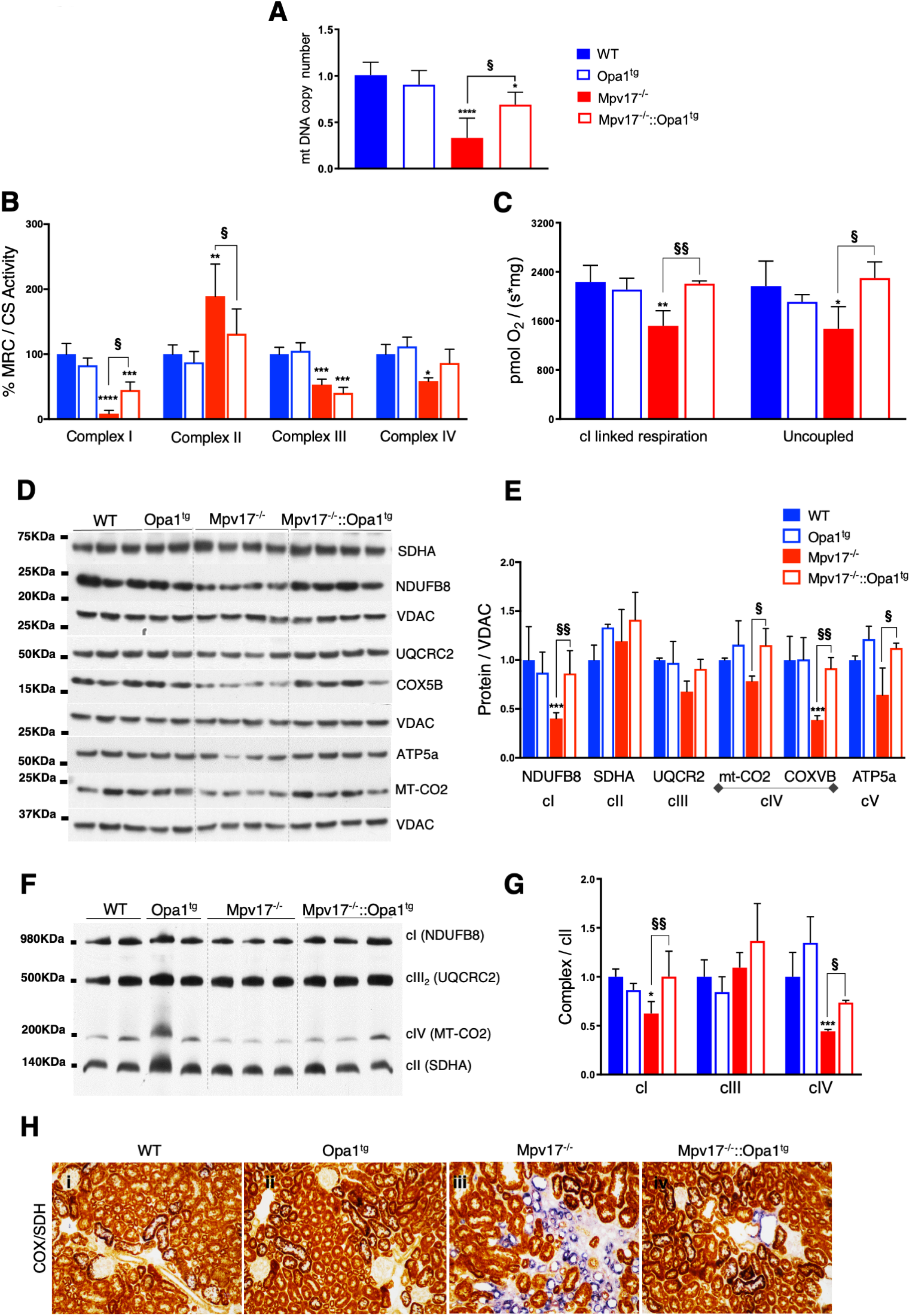
Effects of Opa1 overexpression on mtDNA content and respiratory chain of *Mpv17*^*-/-*^. A) mtDNA copy number in WT, *Opa*^*tg*^, *Mpv17*^*-/-*^ and *Mpv17*^*-/-*^::*Opa1*^*tg*^ kidneys (5 mice/genotype). Symbols * and **§** represent the significance levels of *Mpv17*^*-/-*^::*Opa1*^*tg*^ vs. WT and *Mpv17*^*-/-*^, respectively, calculated by one‐way ANOVA with Tukey’s post hoc multiple comparison test: **** <0.0001 (*Mpv17*^*-/-*^ vs. WT); *p=0.01 (*Mpv17*^*-/-*^::*Opa1*^*tg*^ vs. WT); **§** 0.0137 (*Mpv17*^*-/-*^::*Opa1*^*tg*^ *vs. Mpv17*^*−/−*^). B) MRC activities normalized to citrate synthase (CS) in kidney homogenates of WT, *Opa*^*tg*^, *Mpv17*^*-/-*^ and *Mpv17*^*-/-*^::*Opa1*^*tg*^ mice (n=4 mice/genotype). Symbols * and **§** represent the significance levels of *Mpv17*^*-/-*^::*Opa1*^*tg*^ vs. WT and *Mpv17*^*-/-*^, respectively, calculated by one‐ way ANOVA with Tukey’s post hoc multiple comparison test: CI ****p<0.0001 (*Mpv17*^*-/-*^ vs. WT); *** 0.009 (*Mpv17*^*-/-*^::*Opa1*^*tg*^ vs. WT), **§** p=0.04515 (*Mpv17*^*-/-*^::*Opa1*^*tg*^ *vs. Mpv17*^*−/−*^); CII ** 0.0095 (*Mpv17*^*-/-*^ vs. WT), **§** p=0.0479 (*Mpv17*^*-/-*^::*Opa1*^*tg*^ *vs. Mpv17*^*−/−*^); CIII *** p=0.0005 (*Mpv17*^*-/-*^ vs. WT), *** 0.0002 (*Mpv17*^*-/-*^::*Opa1*^*tg*^ vs. WT); CIV * 0.0121 (*Mpv17*^*-/-*^ vs. WT). C) High resolution respirometry on isolated kidney mitochondria from WT, *Opa*^*tg*^, *Mpv17*^*-/-*^ and *Mpv17*^*-/-*^::*Opa1*^*tg*^ mice (n=4 mice/genotype). Symbols * and **§** represent the significance levels of *Mpv17*^*-/-*^::*Opa1*^*tg*^ vs. WT and *Mpv17*^*-/-*^, respectively, calculated by one‐way ANOVA with Tukey’s post hoc multiple comparison test: CI **\*\*** 0.0032 (*Mpv17*^*-/-*^ vs. WT); **§§** 0.0071 (*Mpv17*^*-/-*^::*Opa1*^*tg*^ *vs. Mpv17*^*−/−*^); uncoupled **\*** p=0.0402 (*Mpv17*^*-/-*^ vs. WT), **§** p=0.0236 (*Mpv17*^*-/-*^::*Opa1*^*tg*^ *vs. Mpv17*^*−/−*^). D) Western-blot immunovisualization of MRC complex subunits in kidney homogenates from the indicated genotypes. E) Densitometric analysis of the Western-blot immunovisualization shown in D (n=3 WT; 3 *Opa1*^*tg*^; 4 *Mpv17*^*-/-*^. Symbols * and **§** represent the significance levels of *Mpv17*^*-/-*^::*Opa1*^*tg*^ vs. WT and *Mpv17*^*-/-*^, respectively, calculated by one‐way ANOVA with Tukey’s post hoc multiple comparison test: CI **\*\*\*** p=0.0007 (*Mpv17*^*-/-*^ vs. WT), **§§** 0.057 (*Mpv17*^*-/-*^::*Opa1*^*tg*^ *vs. Mpv17*^*−/−*^); CIV **§** p=0.04 (*Mpv17*^*-/-*^::*Opa1*^*tg*^ *vs. Mpv17*^*−/−*^); COXVB **\*\*\* p=** 0.0005 (*Mpv17*^*-/-*^ vs. WT), **§§** p=0.003 (*Mpv17*^*-/-*^::*Opa1*^*tg*^ *vs. Mpv17*^*−/−*^); CV **§** p=0.0146 (*Mpv17*^*-/-*^::*Opa1*^*tg*^ *vs. Mpv17*^*−/−*^). F) 1-D BNGE analysis of DDM-solubilized kidney mitochondria from the different genotypes G) Densitometric analysis of the Western-blot immunovisualization shown in in F (n=4 WT, 4 *Opa1*^*tg*^; 7 *Mpv17*^*-/-*^ and *Mpv17*^*-/-*^::*Opa1*^*tg*^. Symbols * and **§** represent the significance levels of *Mpv17*^*-/-*^::*Opa1*^*tg*^ vs. WT and *Mpv17*^*-/-*^, respectively, calculated by two‐way ANOVA with Tukey’s post hoc multiple comparison test: CI * p=0.0218 (*Mpv17*^*-/-*^ vs. WT), §§ p=0.0045 (*Mpv17*^*-/-*^::*Opa1*^*tg*^ *vs. Mpv17*^*−/−*^); CIV*** p=0,0002 (*Mpv17*^*-/-*^ vs. WT), § p=0,0432 (*Mpv17*^*-/-*^::*Opa1*^*tg*^ *vs. Mpv17*^*−/−*^). H) Histochemical staining for COX in kidneys of WT, *Opa*^*tg*^, *Mpv17*^*-/-*^ and *Mpv17*^*-/-*^::*Opa1*^*tg*^ mice. Scale bar: 10μm.

We then analysed the spectrophotometric activities of the MRC complexes in kidney homogenates from animals of the different genotypes. cI and cIV specific activities normalized to citrate synthase (CS) activity, were significantly higher in *Mpv17*^*-/-*^::*Opa1*^*tg*^ *vs. Mpv17*^*−/−*^ samples, although they were still significantly lower than in WT and *Opa1*^*tg*^ animals (Figure 2B). No differences were detected between *Mpv17*^*-/-*^::*Opa1*^*tg*^ *vs. Mpv17*^*−/−*^ in cIII_2_ activity, which was in both strains lower than in control animals. Finally, cII, which was significantly higher in *Mpv17*^*-/-*^ kidneys, was comparable to CTR and *Opa1*^*tg*^ animals in *Mpv17*^*-/-*^::*Opa1*^*tg*^ mice. In addition, cI-dependent (rotenone sensitive) and uncoupled oxygen consumptions, which were both reduced in *Mpv17*^*-/-*^ mice, were normal in *Mpv17*^*-/-*^::*Opa1*^*tg*^ kidney mitochondria (Figure 2C).

Given the above results, we analysed the protein levels of several MRC subunits. In agreement with the spectrophotometric activities, cI subunit NDUFB8, cIV subunits mt-CO2 and COX5B, and cV subunit ATP5A were increased in *Mpv17*^*-/-*^::*Opa1*^*tg*^ *vs. Mpv17*^*−/−*^ mice (Figure 2D, E). No significant changes were detected in cII subunit SDHA and cIII_2_ subunit UQCR2. Finally, we analysed respiratory holocomplexes in dodecyl-maltoside (DDM)-solubilized kidney mitochondria by first-dimension blue native gel electrophoresis (1D-BNGE). The amounts of both cI and cIV, which were decreased in *Mpv17*^*-/-*^ mice, became comparable to CTRs in *Mpv17*^*-/-*^::*Opa1*^*tg*^ (Figure 2F, G). However, Western-blot analyses of 1D-BNGE in digitonin-treated isolated kidney mitochondria did not show any significant changes in the amount of cI- and cIII_2_-containing supercomplexes (SC), suggesting that the amelioration of the phenotype in the *Mpv17*^*-/-*^::*Opa1*^*tg*^ mice was not related to an increase in the stability of MRC SC (Supplementary Figure S4C, D). Histochemical analysis of kidney sections revealed decrease in the intensity of COX-specific staining in virtually all glomeruli, and uneven distribution of COX staining in the tubuli (Figure 2H) of *Mpv17*^*-/-*^, as previously reported (Viscomi *et al*, 2009).

Decreased mtDNA content and expression were previously reported to correlate to mitochondrial coenzyme Q (CoQ) levels (Kühl *et al*, 2017). However, we did not find any difference in CoQ levels in homogenates from *Mpv17*^*-/-*^ and CTR kidneys (Supplementary Figure S4E).

We then used laser-captured microdissected glomeruli and tubuli from animals of the different genotypes. We analysed both COX-positive and COX-negative tubuli from *Mpv17*^*-/-*^ and *Mpv17*^*-/-*^::*Opa1*^*tg*^ individuals. By quantitative PCR, we found that mtDNA content was particularly low in *Mpv17*^*-/-*^ glomeruli and COX-negative tubuli (18% and 29% of the controls, respectively) (Supplementary Figure 5A, B, respectively). In contrast, no differences were detected in COX-positive tubuli compared to CTRs (Supplementary Figure 5C). Notably, the mtDNA content in both glomeruli and tubuli was markedly increased in *Mpv17*^*-/-*^::*Opa1*^*tg*^ *vs. Mpv17*^*−/−*^ mice, although in both cases it remained significantly lower than in WT and *Opa1*^*tg*^ animals (60% and 68% respectively).

These results indicate that *Opa1* overexpression is preventing mtDNA depletion and mitochondria dysfunction in specific regions of *Mpv17*^*-/-*^ mouse kidney.

### *Opa1* overexpression prevents mitochondrial fragmentation and preserves cristae ultrastructure in *Mpv17*^*-/-*^ mice

TEM analysis of the kidneys at low magnification showed a significant reduction in the number of cells in the glomeruli of *Mpv17*^*-/-*^ kidney sections compared to the WT, *Opa1*^*tg*^ and *Mpv17*^*-/-*^::*Opa1*^*tg*^ (Figure 3A, B) with accumulation of abnormal mitochondria with altered cristae in glomerular cells (Figure 3C, D). In addition, a mixed population of normal and altered mitochondria was present in different areas of the tubuli of *Mpv17*^*-/-*^ mice (Supplementary Figure 6).

**Figure 3.**
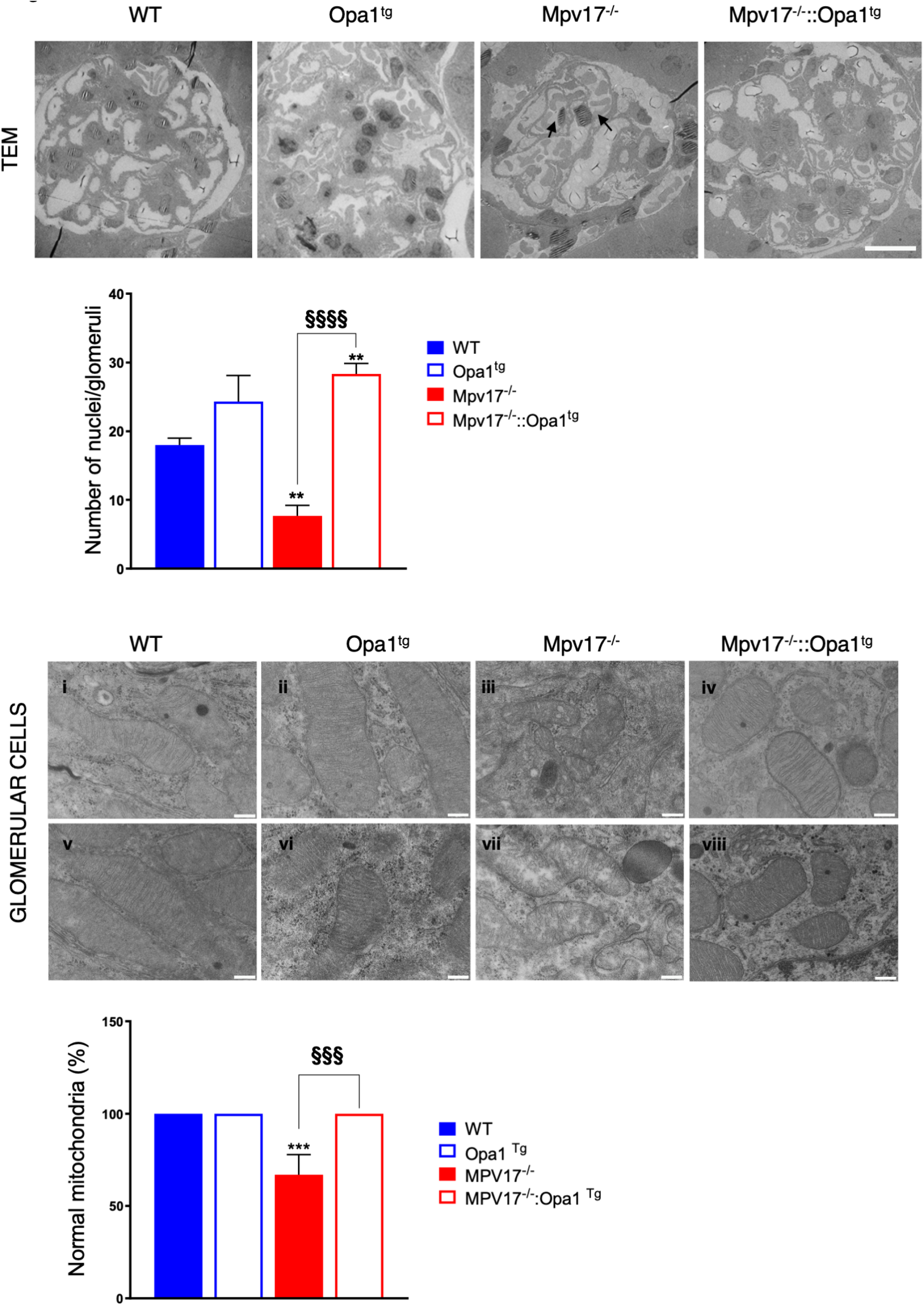
TEM analysis of *Mpv17*^*-/-*^ kidneys. A) Representative low magnification TEM micrographs from mouse kidneys of the indicated genotypes showing the glomeruli. Scale bars: 200 nm. B) Quantification of the number of nuclei/glomeruli in WT, *Mpv17*^*-/-*^ and *Mpv17*^*-/-*^::*Opa1*^*tg*^ kidneys. Data plotted for number of nuclei per glomeruli are mean ± S.D. (n=3). Symbols * and **§** represent the significance levels of *Mpv17*^*-/-*^::*Opa1*^*tg*^ vs. WT and *Mpv17*^*-/-*^, respectively, calculated by one-way ANOVA followed by Tukey’s multiple comparison test:: ** p=0.0021 (*Mpv17*^*-/-*^ vs. WT and *Mpv17*^*-/-*^::*Opa1*^*tg*^ vs. WT); **§§§§** p<0.0001 (*Mpv17*^*-/-*^::*Opa1*^*tg*^ *vs. Mpv17*^*−/−*^). C) Representative TEM images at high magnification showing the accumulation of damaged mitochondria in *Mpv17*^*-/-*^ glomeruli. D) Quantification of the number of damaged mitochondria in glomeruli of the indicated genotypes. Data plotted for number of damaged mitochondria/cell are expressed as mean ± S.DD. (n=3). Symbols * and **§** represent the significance levels of *Mpv17*^*-/-*^::*Opa1*^*tg*^ vs. WT and *Mpv17*^*-/-*^, respectively, calculated by one-way ANOVA followed by Tukey’s multiple comparison test:: *** p=0.0021; * p=0.0403 (*Mpv17*^*-/-*^ vs. WT and *Mpv17*^*-/-*^::*Opa1*^*tg*^ vs. WT); **§§§** p=0.0001; § p=0. 0403 (*Mpv17*^*-/-*^::*Opa1*^*tg*^ *vs. Mpv17*^*−/−*^).

Given the reduced number of cells in *Mpv17*^*-/-*^ glomeruli, we carried out additional experiments in isolated primary podocytes. Since no differences were observed between WT and *Opa1*^*Tg*^ mice we did not produce *Opa1*^*Tg*^ podocytes.

First, we assessed the purity of the culture by immunostaining with an anti-NPHS2 antibody, which recognizes a specific marker for podocytes, and used an anti-cytokeratin antibody to exclude the presence of epithelial cells (Supplementary Figure 7). We then quantified mitochondria length and found a significant decrease in the average size of *Mpv17*^*-/-*^ mitochondria compared to WT, while it was normal in *Mpv17*^*-/-*^::*Opa1*^*tg*^ podocytes (Figure 4 A, B). In keeping with these data, we found that the amount of MFN2, a protein of the outer mitochondrial membrane playing a key role in mitochondrial fusion, was markedly decreased in *Mpv17*^*-/-*^ kidney homogenates, but was comparable to WT in *Mpv17*^*-/-*^::*Opa1*^*tg*^ kidneys (Figure 4C,D). Next, we quantified mitochondrial ultrastructure in isolated podocytes. A significantly reduced number of cristae and increased cristae junction (CJ) width were found in *Mpv17*^*-/-*^ vs. WT cells (Figure 5A-C). In contrast, both number and CJ width were similar to WT in *Mpv17*^*-/-*^::*Opa1*^*tg*^ cells. Similar to what observed for MFN2, MIC19, a component of the MICOS complex involved in cristae shaping (Colina-Tenorio *et al*, 2020), was markedly decreased in *Mpv17*^*-/-*^ kidney mitochondria, but was comparable to WT in *Mpv17*^*-/-*^::*Opa1*^*tg*^ kidneys (Figure 5D,E). In addition, overexposure of the 1D-BNGE membranes revealed accumulation of the free ATP-synthase F1 particle in *Mpv17*^*-/-*^ cells, which was not observed in *Mpv17*^*-/-*^::*Opa1*^*tg*^ (Figure 5F5F). These results suggest that *Opa1* overexpression preserves mitochondrial ultrastructure in the absence of *Mpv17*, possibly by increasing mitochondria fusion and stabilizing MICOS complex and ATP synthase structure.

**Figure 4.**
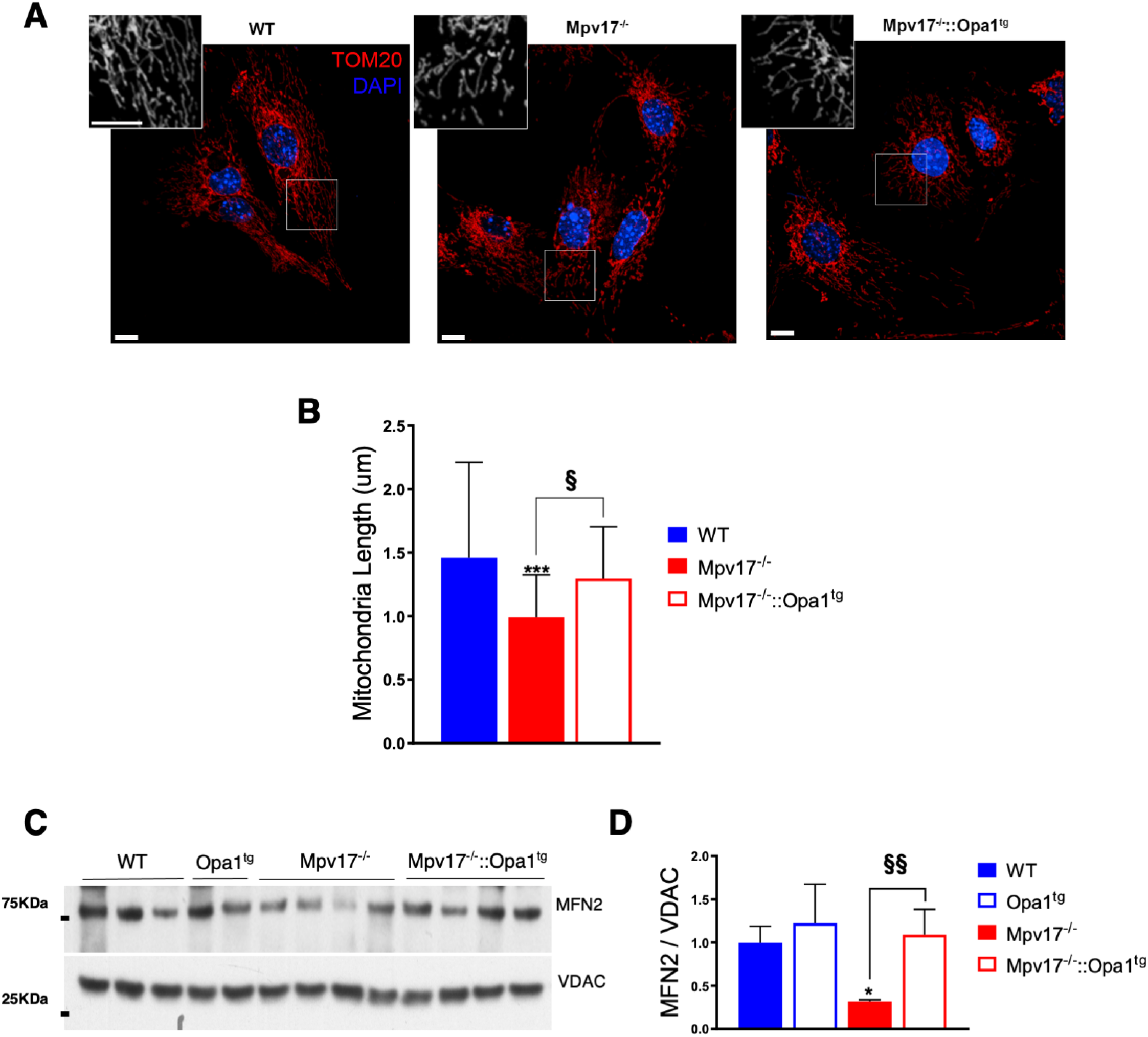
Analysis of isolated podocytes. A) Confocal micrographs showing mitochondrial morphology (TOM20 in red) and nuclei (DAPI in blue) of podocytes from described genotypes. Maximum intensity projection of Z-stacks is shown. Scale bars: 10 μm. B) Mitochondrial length analysis from panel A. Data plotted are mean ± S.E.M. (n=3), total mitochondria (particles) per condition WT= 5663, *Mpv17*^*-/-*^= 5107 and *Mpv17*^*-/-*^::*Opa1*^+/+^= 4360. Symbols * and **§** represent the significance levels of calculated by unpaired t-test: *** p=0.0003 (*Mpv17*^*-/-*^ vs. WT); §§§ p=0.0001 (*Mpv17*^*-/-*^::*Opa1*^*tg*^ *vs. Mpv17*^*−/−*^). C) Western blot immunovisualization with an anti-MFN2 antibody in kidney homogenates. An anti-VDAC antibody was used as a loading control. D) Densitometric analysis of the western blot in C. Symbols * and **§** represent the significance levels of *Mpv17*^*-/-*^::*Opa1*^*tg*^ vs. WT and *Mpv17*^*-/-*^, respectively, calculated by two‐ way ANOVA with Tukey’s post hoc multiple comparison test: CI * p=0.0218 (*Mpv17*^*-/-*^ vs. WT), §§ p=0.0045 (*Mpv17*^*-/-*^::*Opa1*^*tg*^ *vs. Mpv17*^*−/−*^).

**Figure 5.**
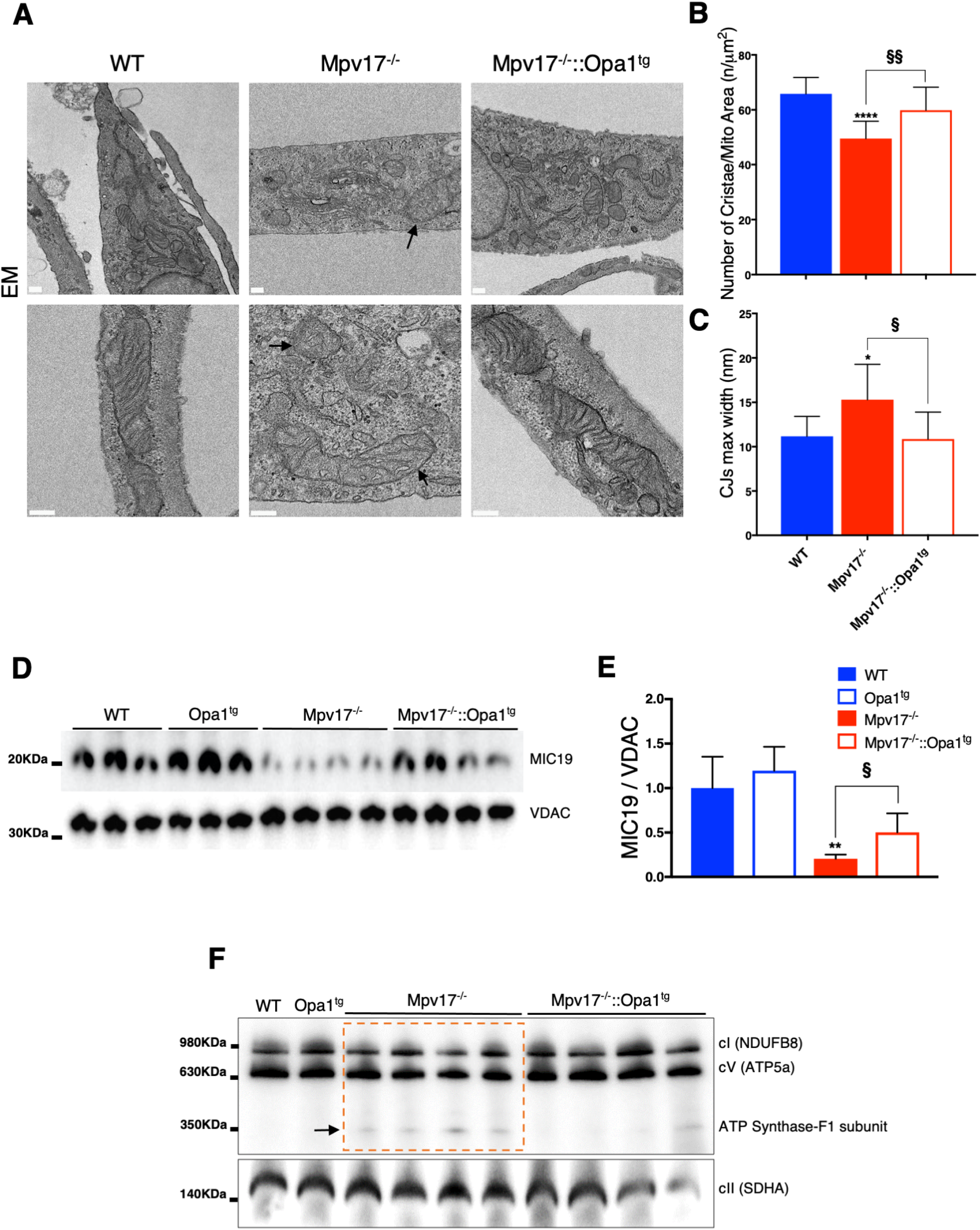
Analysis of mitochondrial cristae morphology. A) Representative TEM images of isolated podocytes for the indicated genotypes. Scale bar: 200 nm. B) Quantification of the number of cristae/mitochondrial area from EM pictures (n= 3/genotype). Data plotted are mean ± S.D. (n=3) of CJs maximum width . Symbols * and **§** represent the significance levels of *Mpv17*^*-/-*^::*Opa1*^*tg*^ vs. WT and *Mpv17*^*-/-*^, respectively, calculated by one‐way ANOVA with Tukey’s post hoc multiple comparison test: ****p<0.0001 (*Mpv17*^*-/-*^ vs. WT), §§ p=0.0068(*Mpv17*^*-/-*^::*Opa1*^*tg*^ *vs. Mpv17*^*−/−*^). C) Quantification of CJ maximum width from EM pictures (n=3). Symbols * and **§** represent the significance levels vs. WT and *Mpv17*^*-/-*^, respectively, calculated by one‐way ANOVA with Tukey’s post hoc multiple comparison test: * p=0.0182 (*Mpv17*^*-/-*^ vs. WT); **§** p=0.0109 (*Mpv17*^*-/-*^::*Opa1*^*tg*^ *vs. Mpv17*^*−/−*^). D) Western-blot immunovisualisation using an anti MFN2 antibody. VDAC was used as a loading control. E) Densitometric analysis of the western blot in D. Error bars represent SD. Symbols * and **§** represent the significance levels of *Mpv17*^*-/-*^::*Opa1*^*tg*^ vs. WT and *Mpv17*^*-/-*^, respectively, calculated by unpaired two‐tailed Student’s t test: ** p=0.0058 (*Mpv17*^*-/-*^ vs. WT); § p=0.0037 (*Mpv17*^*-/-*^::*Opa1*^*tg*^ *vs. Mpv17*^*−/−*^). F) 1D BNGE using anti cI (NDUFB8) and anti cV (ATP5a) antibodies. cII subunit SDHA was used as a loading control. Note the accumulation of ATP synthase F1 subunits in *Mpv17*^*-/-*^ mitochondria.

### *Opa1* overexpression prevents *Mpv17*-dependent glomerulosclerosis by blocking apoptosis

We then investigated whether the correction of mitochondrial ultrastructure can lead to rescue of the glomerulosclerosis and lethality of *Mpv17*^*-/-*^ mice. As mentioned above, a significant reduction in the number of cells in the glomeruli was observed at low magnification in TEM analysis of *Mpv17*^*-/-*^ kidney sections compared to the WT, *Opa*^*tg*^ and *Mpv17*^*-/-*^::*Opa1*^*tg*^ (Figures 33A,B). This result, which suggests increased cell death, prompted us to investigate if *Opa1* overexpression blocked apoptosis in *Mpv17*^*-/-*^ kidneys. Cleaved caspase-3 immunohistochemistry showed a large number of positive cells, especially in the glomeruli of “new” *Mpv17*^*-/-*^ kidneys. In contrast, only a few positive cells were detected in *Mpv17*^*-/-*^::*Opa1*^*tg*^ samples (Figure 6A6A). These results were further confirmed by quantification of cleaved caspase-3 in isolated podocytes (Figure 66B).

**Figure 6.**
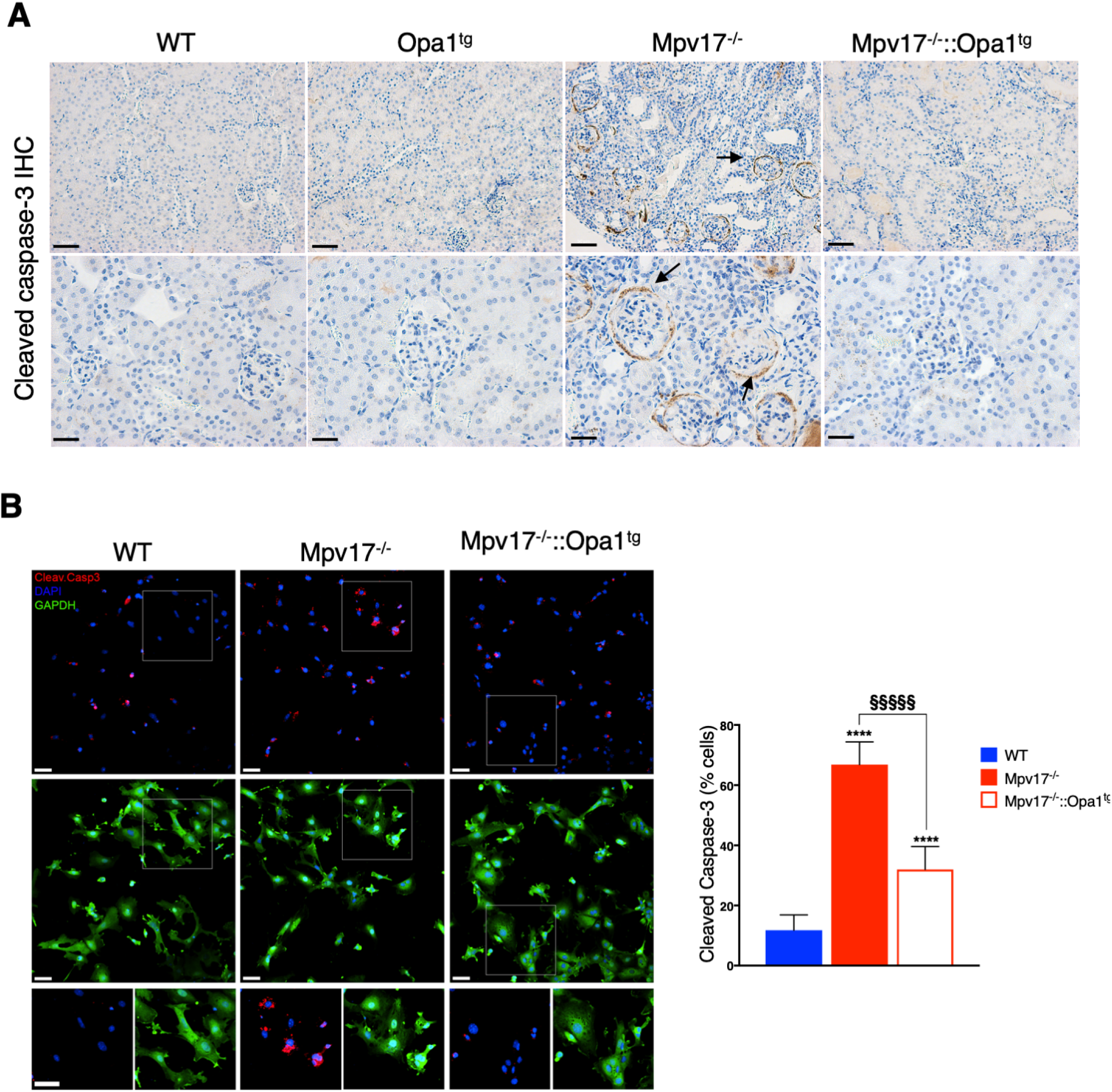
Caspase 3 staining in kidney sections and isolated podocytes. A) A Immunohistochemical staining with anti-caspase 3 antibody. Scale bar: 50μm (upper row); 20μm (lower row). B) Confocal micrographs from primary podocytes derived from described genotypes showing cytoplasm staining (GAPDH in green), apoptotic cells (Cleaved CASPASE-3 in red) and nuclei (DAPI in blue). Maximum intensity projection of Z-stacks is shown. Scale bars: 50 μm. OnOn the right panel, qquantification of caspase 3-positive cells. Data are presented as mean ± SD. Symbols * and **§** represent the significance levels vs. WT and *Mpv17*^*-/-*^, respectively, calculated by one‐way ANOVA with Tukey’s post hoc multiple comparison test^*−*^: **** p<0.0001 (*Mpv17*^*-/-*^ vs. WT and *Mpv17*^*-/-*^::*Opa1*^*tg*^ *vs.* WT); **§§§§** p<0.0001 (*Mpv17*^*-/-*^::*Opa1*^*tg*^ *vs. Mpv17*^*−/−*^).

During apoptosis, the proapoptotic factor Bax translocates from cytoplasm to the mitochondrial outer membrane promoting the release of cytochrome c, which, in turn, interacts with Apaf1 activating the caspase cascade (Figure 7A). Caspase-3 activation also causes the cleavage of the pro-autophagic factor Beclin-1 in three fragments, of 50, 37 and 35 kDa. Cleaved Beclin-1 cannot activate autophagy, and the 50 kDa N-terminal fragment re-localizes to both nucleus and mitochondria, enhancing the apoptotic cascade (Hill *et al*, 2019; Luo & Rubinsztein, 2013; Wirawan *et al*, 2010). We found that Bax expression was increased in *Mpv17*^*-/-*^ mice, whereas its levels were comparable to WT in *Mpv17*^*-/-*^::*Opa1*^*tg*^ littermates (Figure 7B, C). In addition, although several cleaved forms of Beclin-1 were present in WT kidney homogenates, the three previously described, caspase-3 dependent, Beclin-1 fragments were clearly predominant in *Mpv17*^*-/-*^ kidney homogenates. In contrast, the same pattern as in the WT was detected in the *Mpv17*^*-/-*^::*Opa1*^*tg*^ samples (Figure 7D).

**Figure 7.**
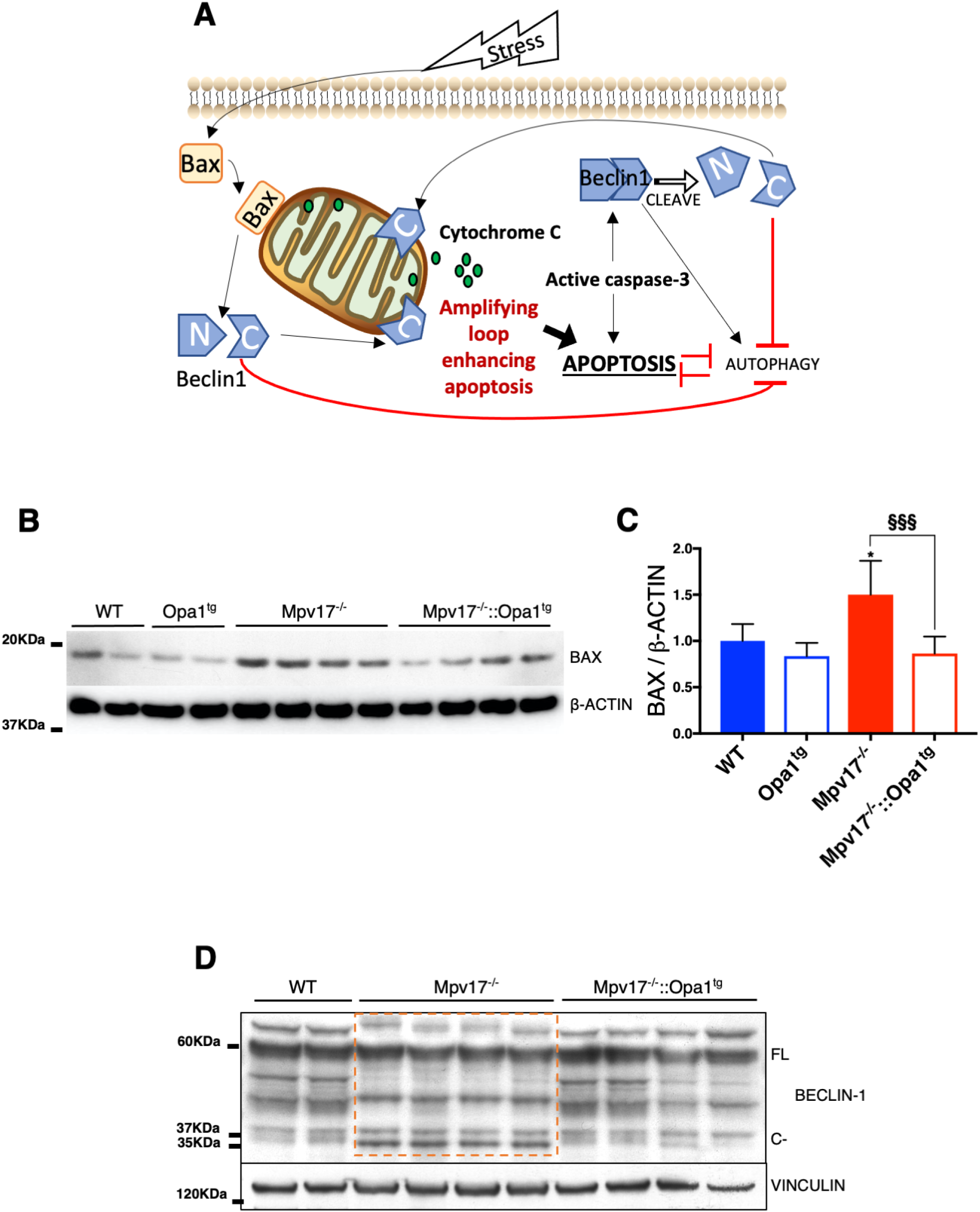
Interplay between apoptosis and autophagy through regulation of Beclin-1. A) Schematic representation of the pathway connecting apoptosis and autophagy via modulation of BECLIN-1 processing. Details in the text. B) Western blot immunovisualization of kidney homogenates using an anti-BAX antibody. C) Densitometric analysis of the western bot in B. Data are presented as mean ± SD (n = 4 WT; n =8 KOS). Symbols * and **§** represent the significance levels vs. WT and *Mpv17*^*-/-*^, respectively, calculated by one‐way ANOVA with Tukey’s post hoc multiple comparison test: * p=0.0176 (*Mpv17*^*-/-*^ vs. WT); §§§ p=0.0008 (*Mpv17*^*-/-*^::*Opa1*^*tg*^ *vs. Mpv17*^*−/−*^). D) Western blot analysis of kidney homogenates using an anti-BECLIN-1 antibody. Note the different pattern of BECLIN-1 processing in Mpv17^*−/−*^ compared to WT and *Mpv17*^*-/-*^::*Opa1*^*tg*^ kidneys.

## Discussion

Alterations in mitochondrial ultrastructure have an impact on mtDNA stability, as demonstrated, for instance, by the presence of multiple deletions in patients carrying mutations in *OPA1* (Hudson *et al*, 2008), *MFN2* (Vielhaber *et al*, 2013) and *ATAD3* (Peralta *et al*, 2018). We showed that moderate overexpression of *Opa1* was able to rescue the mtDNA depletion in “new” *Mpv17*^*-/-*^ kidneys. Although the MPV17 function remains unclear, mutations in its encoding gene constitute a prominent cause of mtDNA instability syndromes in humans. We previously found that mitochondrial cristae ultrastructure was disrupted in the liver of *Mpv17*^*-/-*^ mice, and mtDNA was decreased to less than 10% of the control littermates. However, this was not associated with an overt liver dysfunction unless challenged with ketogenic high fat diet, which led to cirrhosis and mouse liver failure (Bottani *et al*, 2014). We also reported that *Mpv17*^*-/-*^ mice developed a late onset focal segmental glomerulosclerosis, leading to death at around 1.5-2.0 years of age. The early-onset glomerulosclerosis observed after crossing with *Opa1*^*tg*^ mice was unexpected, prompting us to hypothesize the presence of causative variants in one of the supposed genetic modifiers of the *Mpv17*^*-/-*^ mitochondrial phenotypes. The first candidate gene was *Prkdc*, encoding the catalytic subunit of the DNA-dependent protein kinase (DNA-PK), which forms a heterotrimer with the Ku p70/YRCC6-p86/XRCC5 proteins in DNA double-strand break repair and recombination. A spontaneous mutation R2140C in *Prkdc* was found in some mouse strains leading to profound decrease in the amount of the protein, which was proposed to modify the severity of *Mpv17* kidney disease (Papeta *et al*, 2010). However, our mice did not carry the *Prdkc* hypomorphic mutant allele.

A second potential modifier of the *Mpv17*^*-/-*^ phenotype was the nicotinamide nucleotide transhydrogenase (*Nnt*), which regulates NAD(P)H levels by coupling hydride transfer between NAD(H) and NADP(+) to proton translocation across the IMM (Kampjut & Sazanov, 2019), thus regulating redox poise. C57Bl/6J mice carry two spontaneous mutations in the *Nnt* gene, a missense (methionine to threonine) mutation in the mitochondrial leader sequence; and an in-frame deletion that removes exons 7-11, resulting in expression of a truncated, non-functional protein. The mutant Nnt allele was previously proposed to impact on the severity of mtDNA mutations (McManus *et al*, 2019). However, although we detected both the wild-type and truncated forms of the *Nnt* gene in our colony, these variants did not segregate with the disease. These results conclusively suggest that other modifiers may be responsible for the shift in the phenotype. RNAseq analysis showed a few quantitative differences between the “old” KO and the “new” KO strains, including only two mitochondrial targeted gene products, an unknown transporter (*Slc25a48*) and uridine phosphorylase 2. These observations, as well as the few hundred SNP variants identified in the same RNAseq experiment, warrant validation and further investigation in the near future. Mitochondrial cristae are controlled by a molecular network formed by the mitochondrial contact site and cristae organizing system (MICOS), the F_1_F_0_-ATP synthase, the IMM pleiotropic OPA1 protein, and the non‐bilayer‐forming phospholipids cardiolipin and phosphatidylethanolamine (Colina-Tenorio *et al*, 2020). OPA1 has been shown to be epistatic to MICOS (Glytsou *et al*, 2016), and that its protective effect against mitochondrial dysfunction is mediated by ATP synthase oligomerization (Quintana-Cabrera *et al*, 2018). We found that *Opa1* overexpression was able to correct mitochondrial ultrastructure and increase mtDNA content in the kidney of “new” *Mpv17*^*-/-*^ mice. However, whilst the effect on liver mtDNA was modest, the mtDNA increase in kidney was sufficient to preserve the activities of MRC complexes to WT levels. Mechanistically, the correction of mitochondrial ultrastructure was likely mediated by ATP synthase stabilization, as shown by our 1D-BNGE Western-blot data, showing the accumulation of free F_1_ ATP-synthase particles in *Mpv17*^*-/-*^, which were absent in *Mpv17*^*-/-*^::*Opa1*^*tg*^ kidneys. These results are also in agreement with recent published data (Quintana-Cabrera *et al*, 2018). The normalization of mitochondrial ultrastructure explains the reduced apoptosis in *Mpv17*^*-/-*^::*Opa1*^*tg*^ vs. *Mpv17*^*-/-*^ kidneys and podocytes. This process may be boosted by the block of Beclin-1 processing.

Overall, our results further support the deleterious role of cristae disruption in the pathogenetic mechanisms leading to organ failure in mitochondrial disorders, and the correcting effects of moderate overexpression of *Opa1*, which can be exploited for future therapeutic approaches.

## Material and methods

### RNA sequencing analysis

RNA was extracted by a standard Trizol-based protocol from kidney tissue. RNA seq was carried out by the NGS facility, Department of Pathology, University of Cambridge, Cambridge UK. A detailed description of the methodology followed for the collection and analysis of data is reported in the Supplementary Material.

### Antibodies

Mouse monoclonal anti-GAPDH (1:1,000; #ab8245; Abcam), rabbit monoclonal anti-TOM20 Alexa Fluor 647 (1:500; #ab209606; Abcam), Alexa Fluor 488 Goat anti-mouse (1:300; #A11004; Invitrogen), Alexa Fluor 647 Goat anti-rabbit (1:300; #A27040; Invitrogen), rabbit polyclonal anti-NPHS2 (1:300; #ab50339; Abcam), mouse monoclonal anti-CYTOKERATIN (1:300; #ab86734; Abcam), anti-cleaved caspase 3 (1:200; Cell Signalling, #9661).

### Chemicals

All chemicals were from Merck-Sigma Aldrich.

### Animals

All animal experiments were carried out in accordance with the UK Animals (Scientific Procedures) Act 1986 (PPL: P6C20975A) and EU Directive 2010/63/EU. The mice were kept on a C57Bl6 background, and WT littermates were used as controls. The animals were maintained in a temperature- and humidity-controlled animal-care facility with a 12-hr light/dark cycle and free access to water and food and were sacrificed by cervical dislocation.

### Isolation of primary podocytes

Kidneys were removed from 7-to 10-day-old mice, washed in Ca^2^ and Mg^2^ free Hanks medium (Thermo Fisher), and treated with collagenase type I AS (Sigma Aldrich) 1.5 mg/ml, for 1 min at 37°C. The reaction was stopped by growth medium consisting of Dulbecco’s modified Eagle medium: F12 (Thermo Fisher) supplemented with 10% FBS (Gibco), 1x Insulin-Transferrin-Selenium-Ethanolamine (Thermo Fisher), 1 uM hydrocortisone (Sigma Aldrich), 1x Penicillin-Streptomycin and 2 mM L-glutamine (Sigma-Aldrich).

Glomeruli were isolated by sieving through the 100-μm mesh (Fisherbrand™) and seeded in 100 mm dishes precoated overnight at 4 °C with collagen type IV (Sigma-Aldrich). Cells were kept at 37°C in 5% CO_2_ atmosphere for 7 days. Additional sieving was performed trough the 40-μm mesh resuspending the cells with trypsin-EDTA. Cells were plated on dishes or coverslips treated with Poly-L-Lysine (Sigma-Aldrich) and the next day fixed for the respective technique. Cell characterization was performed by immunofluorescence using the podocyte (NPHS2) and epithelial cells (CYTOKERATIN) markers.

### Immunofluorescence and imaging

Cells were fixed with 3.7% formaldehyde in F-12 complete media, permeabilized with 0.1% in PBS with 0.1% Triton X-100 and 0.05% sodium deoxycholate, before staining with primary and secondary antibodies in blocking solution (5% goat serum). Coverslips were mounted with ProLong Glass with or without DAPI (Invitrogen).

Images were acquired randomly for each slide using a Dragonfly Spinning Disk imaging system (Andor Technologies Ltd.), composed by a Nikon Ti-E microscope, Nikon 100x TIRF ApoPlan or 20x ApoPlan objective and an Andor Ixon EMCCD or Zyla sCMOS camera. The z-stacks were acquired using Fusion software (Andor Technologies) and the 3D images analysed and exported using Imaris software (Bitplane Inc.). For mitochondria length measurements 3 preparation of podocytes (n=3) per group was used and several regions of interests (ROIs) were drawn in the peripheric regions of the mitochondrial network using ImageJ (Schindelin *et al*, 2012). Automatic threshold was used and “particles” (mitochondria) bigger than 0.05 μm^2^ were analysed. Cleaved CASPASE-3 analysis was performed counting manually the number of cells (DAPI+GAPDH positive) and CASPASE-3 positive cells (n=3).

### Transmission electron microscopy (TEM)

For TEM analysis, mice were subjected to perfusion with the fixative solution (2.5% Glutaraldehyde; Sigma-Aldrich, 2% Paraformaldehyde; MP Biomedicals in 0.1 M PB buffer; Sigma-Aldrich), 1 mm^3^ pieces of kidney were kept in fixative solution overnight and samples were washed several times in 0.1 M phosphate buffer the following days. For fibroblasts, 1×10^6^ fibroblasts were fixed in the fixative solution for 1 h at RT in agitation. Cells were scraped, centrifuged (2,500 RPM for 10 min at 4°C) and washed 3 times with 0.1 M PB buffer. Pelleted cells were sent to the Electron Microscopy Platform (Scientific and Technological Centers, University of Barcelona, Spain) for sample preparation. Regions containing the glomeruli were selected and ultrathin sections (55 nm) were cut, for kidney and cells, and mounted on 200 mesh copper grids with carbon supported film. Image acquisition (Digital Micrograph, v. 1.85.1535, Gatan, USA) was performed with a transmission electron microscope (FEI Tecnai G2 Spirit Twin) coupled with an CCD camera (Orius-SC200B, Gatan, USA). Quantification of number and width of cristae was performed manually using ImageJ, and 10 cells were analysed for eachgenotype (n=3) and number of cristae was counted per visible mitochondria area, maximum cristae junction (CJ) width was determined by measuring randomly 3-4 CJs/mitochondria. Foot process was measured by the ratio between maximum height and width touching the glomerular basement membrane. The mean of this ratio (Foot Process Aspect Ratio) was calculated from 21 randomly chosen segments per group (n=3).

### Immunoblotting

Mouse tissues were homogenized in ten volumes of 10 mM potassium phosphate buffer (pH 7.4). Mitochondrial-enriched fractions were collected after centrifugation at 800g for 10 min in the presence of protease inhibitors, and frozen and thawed three times in liquid nitrogen. Protein concentration was determined by the Lowry method. Aliquots, 30 μg each, were run through a 12% SDS-PAGE and electroblotted onto a PVDF membrane, which was then immunodecorated with different antibodies.

For BNGE analysis, brain mitochondria isolated as previously described(Fernández-Vizarra *et al*, 2002), were resuspended in 1.5 M aminocaproic acid, 50 mM Bis-Tris-HCl pH 7 and 4 mg dodecylmaltoside /mg of proteins and incubated for 5 min on ice before centrifuging at 20,000 3 g at 4° C. 5% Coomassie G250 was added to the supernatant. 100 μg were separated by 4%–12% gradient BNGE and electroblotted on nitrocellulose membranes for immunodetection.

### Spectrophotometric analysis of respiratory chain activities

Tissues were snap-frozen in liquid nitrogen and homogenized in 10 mM phosphate buffer (pH 7.4). The spectrophotometric activity of cI, and CS was measured as previously described

### Morphological analysis

For histological and immunohistochemical analyses animals were fixed in 10% neutral buffered formalin (NBF) for a few days at room temperature and then included in paraffin wax. 4 μm-thick sections were used for analysis. Haematoxylin-eosin (H&E) staining was performed by standard methods. Immunohistochemistry was performed using a Novolink Polymer Detection System and specific antibodies against the indicated proteins.

### Statistical analysis

All numerical data are expressed as mean ± SD. One- or two-way ANOVA tests with Tukey correction were used for multiple comparisons (see figure legends for details); Kaplan-Meier distribution and log-rank test were used for survival probability analysis. Differences were considered statistically significant for p < 0.05.

## Acknowledgements

Our research was supported by the Telethon Foundation (grant GGP19007), ERC Advanced Grant FP7‐3222424 and NRJ‐Institut de France Grant (to M.Z.); Associazione Luigi Comini ONLUS; and core grant from the Medical Research Council (Grant MC_UU_00015/5). We thank the MRC-Laboratory of Molecular Biology Electron Microscopy Facility for access to their resources.

